# Minerva: An Alignment and Reference Free Approach to Deconvolve Linked-Reads for Metagenomics

**DOI:** 10.1101/217869

**Authors:** David C. Danko, Dmitry Meleshko, Daniela Bezdan, Christopher Mason, Iman Hajirasouliha

## Abstract

Emerging Linked-Read technologies (aka Read-Cloud or barcoded short-reads) have revived interest in standard short-read technology as a viable way to understand large-scale structure in genomes and metagenomes. Linked-Read technologies, such as the 10X Chromium system, use a microfluidic system and a set of specially designed 3’ barcodes (aka UIDs) to tag short DNA reads which were originally sourced from the same long fragment of DNA; subsequently, these specially barcoded reads are sequenced on standard short read platforms. This approach results in interesting compromises. Each long fragment of DNA is covered only sparsely by short reads, no information about the relative ordering of reads from the same fragment is preserved, and typically each 3’ barcode matches reads from 2-20 long fragments of DNA. However, compared to long read platforms like those produced by Pacific Biosciences and Oxford Nanopore the cost per base to sequence is far lower, far less input DNA is required, and the per base error rate is that of Illumina short-reads.

The use of Linked-Reads presents a new set of algorithmic challenges. In this paper, we formally describe one particular issue common to all applications of Linked-Read technology: the deconvolution of reads with a single 3’ barcode into clusters that correspond to a single long fragment of DNA. We introduce **Minerva**, A graph-based algorithm that approximately solves the barcode deconvolution problem for metagenomic data (where reference genomes may be incomplete or unavailable). Additionally, we develop two demonstrations where the deconvolution of barcoded reads improves downstream results: improving the specificity of taxonomic assignments, and by improving clustering of related sequences. To the best of our knowledge, we are the first to address the problem of barcode deconvolution in metagenomics.

## 1 Introduction

### 1.1 Linked-Read Sequencing

Recently, long-read sequencing technologies (e.g. PacBio, Oxford Nanopore) have become commercially available. These techniques promise the ability to improve *de novo* assembly, particularly in metagenomics [5, 9]. While these technologies offer much longer reads than standard short-read sequencing, their base-pair error rates are substantially higher than short reads (10-15% error vs. 0.3%). More important, long-read technologies have substantially higher costs, lower throughput, and require large amounts of DNA, or PCR amplification, as input. Currently, this makes long-reads impractical for large-scale screening of whole genome or metagenome samples and most low-input clinical samples.

As an alternative, low-cost, and low-input (~1 ng) DNA library preparation techniques using microfluidic 3’ barcoding methods have recently emerged (e.g. Moleculo/Illumina, 10X Genomics) that address these shortcomings. With these new technologies, input DNA is sheared into long fragments of ~10-100kbp. After shearing, a 3’ barcode is ligated to short reads from the fragments such that short reads from the same fragment share the same 3’ barcode (N.B. the 3’ barcode is unrelated to the standard 5’ barcode used for sample multiplexing). Finally, the short reads are sequenced using industry standard sequencing technologies (e.g. Illumina Hiseq). This process is commonly referred to as, Linked-Read sequencing. Linked-Reads offer additional *long-range* information over standard short-reads.

Reads with matching 3’ barcodes are more likely to have emerged from the same fragment of DNA than two randomly sampled reads. However, each fragment of DNA is only fractionally covered by reads. This increases the amount of long range information obtained from a given experiment but makes it impossible to assemble reads from a single barcode into a contiguous stretch of sequence. This tradeoff has been used recently to phase large-scale somatic structural variations [7, 17].

State of the art Linked-Read sequencing systems use the same 3’ barcode to label reads from several fragments of DNA. Existing systems have a relatively limited number of 3’ barcodes; loading multiple fragments of DNA into the same 3’ barcodes is critical for high throughput experiments. In particular, in our work using the 10X Genomics system, we observed that there were 2-20 long fragments of DNA per 3’ barcode and that 3’ barcodes with more reads tended to have more fragments. This can complicate downstream applications. In the absence of other information, it is difficult to distinguish the random assortment of reads into a 3’ barcode from an actual structural variation or a different source genome.

To address this critical issue, we define the barcode deconvolution problem. Briefly, each group of reads that share a 3’ barcode has an unobserved set of fragments from which each read was drawn. The barcode deconvolution problem is the problem of assigning each read with a given 3’ barcode to a group such that every read in the group came from the same fragment and so there is only one group per fragment. We note that a fragment assignment is stricter than genomic assignment. Each read from the same fragment necessarily came from the same genome but it is possible to have multiple fragments from the same genome whose reads share the same 3’ barcode.

Linked-Reads provide significantly more information about the proximity of short reads than standard short read sequencing. With the exception of very common or repetitive sequences, the co-occurrence of particular sequences across several 3’ barcodes provides evidence that the co-occurring sequences were drawn from the same underlying DNA molecules.

### 1.2 Glossary

We define several of the terms we use throughout this paper:

**Fragment** A long (10-100kbp) piece of DNA.
**3’ Barcode** The barcode sequence that tags reads from the same fragment of DNA.
**Read Cloud** All reads that share a single 3’ Barcode, even if the reads emerged from different fragments.
**Enhanced Read Cloud** A set of reads from a single sample that all included the same 3’ barcode sequence and that were grouped together by a solution to the barcode deconvolution problem.

### 1.3 Applications of Linked-Reads in Metagenomics

Linked-Reads have several potential benefits for metagenomic research compared to standard short-reads. Linked-Reads carry information about long stretches of sequence. In principle this information can be used to improve taxonomic classification of reads, improve the assembly of microbial genomes, identify horizontally transferred sequences, quantify the genetic structure of low-abundance organisms and catalog intra-sample genetic structural variants. In the near term, algorithms for analyzing short read sequences can be used on Linked-Read data without modification which makes linked reads a practical choice for many studies.

Compared to long-read sequencing, Linked-Reads can be used to sequence samples far more deeply for the same amount of money and can accept much smaller amounts of input DNA. This is important for metagenomics; even at the same read depth, Linked-Reads may be more useful for studying low abundance organisms because Linked-Reads span a much longer stretch of a genome for the number of bases sequenced (i.e. very high physical coverage) and could be used to resolve microbial structural variation.

In this paper, we address the barcode deconvolution problem, a fundamental problem of using Linked-Reads for metagenomics. We show that addressing the barcode deconvolution problem improves downstream results for two demonstration applications.

### 1.4 The Barcode Deconvolution Problem

We formally define the barcode deconvolution problem for a single 3’ barcode. We note that our solution requires a large number of 3’ barcodes but this is not necessary to state the problem generally.

As input we are given a set of *n* reads from the same Read Cloud. Each read has the same 3’ barcode and an unobserved class that represents the fragment from which the read was drawn. For a given 3’ barcode with *n* reads drawn from *f* fragments we have 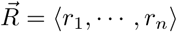 where *r_i_* represents the unobserved class for read *i*, and 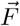, the set of possible fragments 〈*f*_1_, · · ·, *f_n_*〉.

A solution to the barcode deconvolution problem for a single read cloud would be a function mapping a set of read classes to fragment classes

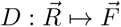

such that the function produces the same value for reads from the same fragment for all *n* reads in *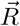*

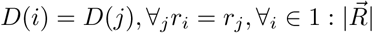

For a set of Read Clouds a solution to the barcode deconvolution problem would be the set of solutions for each individual Read Cloud.

When a reference genome is available the barcode deconvolution problem is relatively trivial, so long as major structural variants are absent. All individual reads from the same Read Cloud can be mapped to the reference genome using any good read alignment method. Reads from the same fragment will tend to be clustered near one another on the reference genome; there is little chance that reads from different fragments in the same Read Cloud will be proximal, unless the reference genome is very small. However, if a reference genome is not available, is small, or structural variation is present read mapping may not provide a good solution to the barcode deconvolution problem. All of these conditions are common in metagenomics.

To the best of our knowledge, we are the first to formally describe the barcode deconvolution problem.

### 1.5 Summary

We have developed a novel method, **Minerva**, that explicitly uses information from sequence overlap between Read Clouds to approximately solve the barcode deconvolution problem for metagenomic samples. Our approach was inspired by topic modeling in Natural Language Processing (NLP) which studies methods to find groups of co-occurring words in text. We demonstrate how our technique can be effectively applied to real metagenomic Linked-Read data (section 2.4) and improve analysis (sections 2.5, 2.6) for two example use cases.

We present our solution to the barcode deconvolution problem in details (section 4.2). We also develop a probabilistic generative model justifying key assumptions of our procedure (section 4.1). We report our negative results, models that we tested but performed poorly (section 7.4).

## 2 Results

### 2.1 Algorithm Overview

We have developed **Minerva**, an algorithm that approximately solves the barcode deconvolution problem for metagenomics. Minerva works by matching reads from the same Read Cloud that share kmers with reads from other Read Clouds. This algorithm processes each Read Cloud individually by building a sparse graph between reads and other Read Clouds, converting that graph into a graph between reads, and clustering that graph. This method is discussed in detail in 4.2.1.

### 2.2 Primary Data Sets

We tested Minerva using two primary real data sets from two microbial mock communities. The first community (Dataset 1) contained five bacterial species: *E. coli, Enterobacter cloacae, Micrococcus luteus, Pseudomonas antarctica,* and *Staph. epidermidis.* The second community (Dataset 2) contained 8 bacterial species and 2 fungi: *Bacillus subtilis, Cryptococcus neoformans, Enterococcus faecalis, E. coli, Lactobacillus fermentum, Listeria monocytogenes, Psuedomonas aeruginosa, Saccharomyces cerevisiae, Salmonella enterica*, and *Staphylococcus aureus*.

We elected to use mock communities over simulated data as we are not aware of any tools that simulate Linked-Read data for metagenomic communities. The mock communities chosen are standard microbial positive controls as noted by [11].

Roughly 1ng of high molecular weight (HMW) DNA was extracted from each sample. The HMW DNA was processed using a 10X Chromium instrument and we prepared sequencing libraries. Each library was sequenced on an Illumina Hiseq with 2×150 paired-end reads. Roughly 20M reads were generated for each sample, for testing we selected 10M reads from each while ensuring that we only selected complete barcodes. Both samples showed some evidence of human contamination, reads that mapped to the human genome were not removed from the samples (but were not used to generate statistics on barcode purity) since some amount of human DNA is typical in metagenomic samples.

In both samples reads were distributed over 3*10^6^ barcodes. We used *LongRanger Basic* [1] to attach barcodes to reads and perform error correction on barcodes. Both samples has a similar number of reads per barcode. Sample 2 had more species represented in each barcode on average, though not necessarily more fragments since fragments can originate from the same genome. Statistics about the datasets are summarized in table 1. Both datasets are available through the 10X Metagenomics Consortium and on our project GitHub page.

**Table 1:** Dataset Properties

|  | Dataset 1 | Dataset 2 |
| --- | --- | --- |
| Number of Read Pairs | $10^7$ | $10^7$ |
| Number of Species | 5 | 10 |
| Mean Read Cloud Richness | 2.74 | 5.79 |
| Mean Read Cloud Size | 7.399 | 7.515 |
| Barcode N50 Size | 11 | 11 |
| Barcode N90 Size | 4 | 4 |

We determined the actual fragment of origin for each read by mapping reads to the source genomes and clustering positions in case multiple fragments from the same genome were present in the same read cloud.

### 2.3 Runtime and Performance

Minerva’s runtime performance largely depends on two parameters: *K,* the size of the kmers used to match reads and Anchor Dropout, the minimum size the Read Cloud being deconvolved. We list the total runtime and RAM usage for Minerva (table 2) on both of our test datasets with different parameters. We note that our implementation of Minerva is single threaded but that the algorithm itself is trivially parallelizable.

**Table 2:** Runtime Performance

| Dataset | K | Anchor Dropout | Runtime (hours) | RAM (GB) |
| --- | --- | --- | --- | --- |
| Dataset 1 | 20 | 50 | 1.48 | 93 |
| Dataset 1 | 30 | 50 | 1.71 | 163 |
| Dataset 2 | 20 | 50 | 3.27 | 115 |
| Dataset 2 | 30 | 50 | 3.66 | 200 |
| Dataset 1 | 20 | 30 | 12.84 | 92 |
| Dataset 1 | 30 | 30 | 15.5 | 163 |
| Dataset 2 | 20 | 30 | 36.69 | 115 |
| Dataset 2 | 30 | 30 | 40 | 200 |

### 2.4 Minerva Approximately solves the Barcode Deconvolution Problem

Minerva was able to identify subgroups in Read Clouds that largely corresponded to individual fragments of DNA. We term these subgroups ‘Enhanced Read Clouds’. We measured the quality of each enhanced Read Cloud using two metrics: shannon entropy index 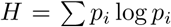 and purity 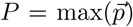 where *p_i_* indicates the proportion of an enhanced Read Cloud that belongs to each fragment. These values are shown in figure 2 as compared to Read Clouds which were not enhanced (’standard’ Read Clouds). In general, Minerva produced a large number of perfect (*P* = 1, *H* = 0) enhanced Read Clouds.

**Figure 1:**
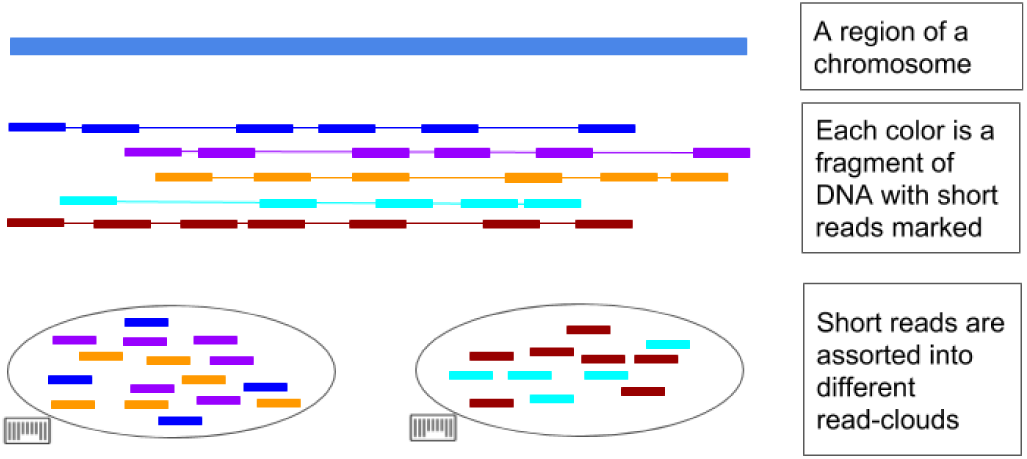
Schematic representing Linked-Read sequencing. Different colors represent reads from different fragments that can be mixed together and tagged with the same barcode.

**Figure 2:**
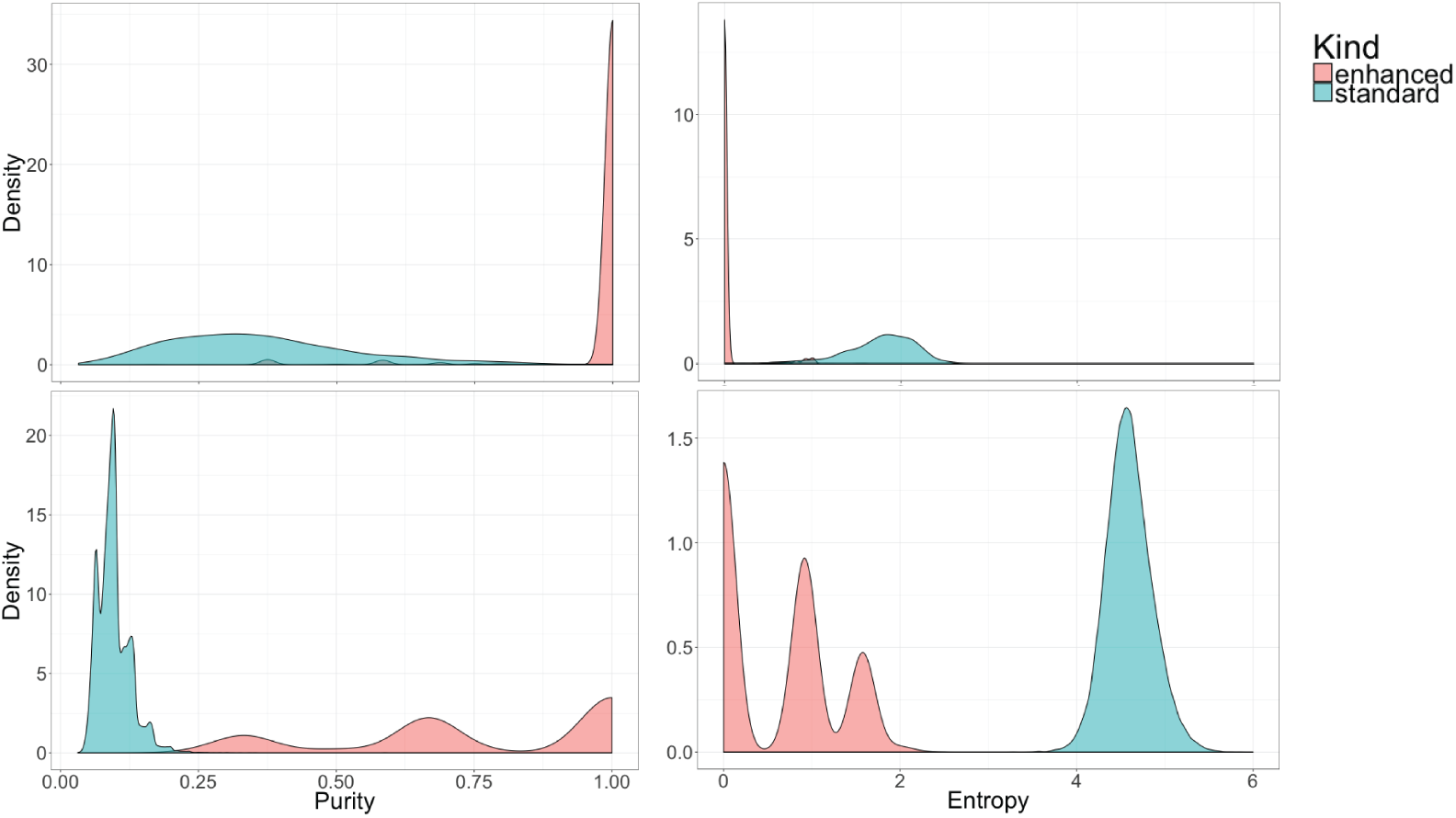
Clockwise from top left 1) Purity in dataset one for enhanced and 3’ barcodes 2) Shannon index in dataset one for enhanced and 3’ barcodes 3) Shannon index in dataset two for enhanced and 3’ barcodes 4) Purity in dataset two for enhanced and 3’ barcodes.

We also tested whether the quality of the enhanced Read Clouds changed with the number of reads in the Read Cloud. We found a small inverse relationship between Read Cloud size and purity but established that our previous results were not being inflated by a large number of very small enhanced Read Cloud (figure 7). Note that enhanced Read Clouds of size 1 would be trivially perfect and are always excluded from results.

In testing we found three parameters that seemed to have the most effect of Minerva’s performance. The number of links required between reads to form a cluster (*eps*), the kmer size used to make minimizing kmers (*K*), and the maximum allowed frequency of each read *(maxk).* In supplementary figure 1 we show how these parameters affect Minerva’s performance under three different metrics: mean enhanced barcode purity, mean enhanced barcode size, and total reads clustered. Large, pure, and complete clusters being the ideal. We found that Minerva’s parameters could be used to tune performance between very large and very pure enhanced barcodes depending on the downstream application.

### 2.5 Enhanced Read Clouds can be Clustered into Meaningful Groups

After deconvolving barcodes into Enhanced Read Clouds it is useful to group Enhanced Read Clouds that likely came from the same genome. This is essentially a clustering problem. Initially we explored graph based approaches similar to our algorithm for Read Cloud deconvolution. These algorithms relied on the assumption that elements being clustered would have small numbers of distinguishing elements and a relatively high *a priori* probability of originating from the same cluster. When dealing with individual barcodes these assumptions proved reasonable; faced with the complexity of a full dataset these assumptions became inaccurate and graph based algorithms performed poorly.

With relatively little structure in the data that could be known *a priori* we turned to topic modeling algorithms to discover implicit genetic structures in our data. Latent Dirichlet Allocation is a classic model in Natural Language Processing [3]. LDA is a generative model that assumes data was created using a certain, well defined, stochastic process. Unlike our graph based algorithm this model had a large number of parameters that could be adjusted until a model that fit the data well was found. Training the model consists of finding parameters that make it more likely that the observed data would be generated using the given stochastic process; typically this is done with Gibbs sampling.

Typically LDA is used to analyze corpora of natural language. Natural language corpora are organized into documents (e.g. emails or book chapters) that consist of words. The base version of LDA does not consider what order words in a document occur, just how often each word occurs in a given document; this is referred to as a bag-of-words model. Formally, documents are modeled as a sparse vector over a large vocabulary of words where entries represent the number of times a word occurs in the document. LDA maps documents from a high dimensional word-space to a lower dimensional topic-space. In NLP topics typically have intuitive interpretations as thematically consistent units. A key advantage of LDA is that it can classify words based on context (i.e. a river bank vs. a financial bank), this may be useful for classifying conserved motifs.

We used LDA to cluster Read Clouds (represented as sets of kmers). Each topic generated by LDA was considered to be a single cluster.

We used LDA to project enhanced and standard Read Clouds into a lower dimensional space. We treated each Read Cloud as a document containing minimum sparse kmers as words. We removed kmers that occured far more often than average in a process similar to removing stop-words in NLP. We ran LDA with hyper-parameter optimization on our Read Cloud-documents and clustered to obtain a topic vector for each Read Cloud using the implementation LDA in Mallet [12]. Using X-Means we clustered the topic vectors representing Read Clouds into discrete groups.

With standard Read Clouds LDA essentially cannot distinguish any structure, with enhanced Read Clouds LDA can generate clusters that are less diverse. The clusterings can be compared in figure 3. This could be used to improve assemblies by clustering similar reads and reducing spurious connections. Note that we denote chromosomes rather than genomes in the figure since our process does not attempt to link chromosomes from the same organisms.

**Figure 3:**
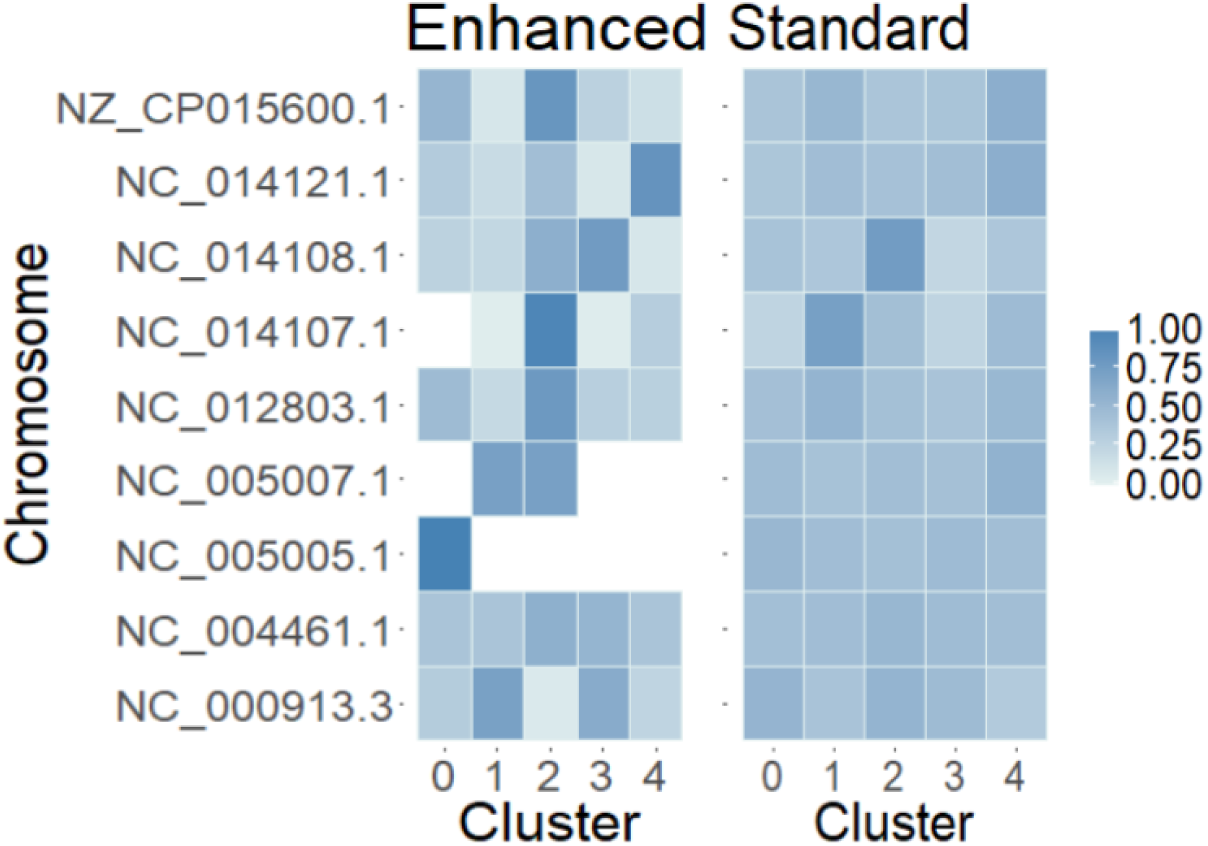
Abundance of different chromosomes across clusters as assigned by LDA. Enhanced Read Clouds dramatically improve LDA’s ability to distinguish structure in Dataset 1. This figure uses the same deconvolution as figure 2

### 2.6 Enhanced Read Clouds Improve Short Read Taxonomic Assignment

We observed that reads from a single linked fragment could be classified using any short read taxonomic classifier. These classifiers, however, often have trade offs between recall and precision. Enhanced Read Clouds can be used to improve recall of a short read classifier without harming precision.

Many of the reads classified by short read taxonomic classifiers cannot be assigned to low taxonomic ranks. However, all reads from the same fragment of DNA must all have the same taxonomic rank. Read Clouds can be used to promote unspecific taxonomic assignments. Any read with a taxonomic rank that is an antecedent of a lower taxonomic rank in the same Read Cloud can be promoted to the lower rank, provided there are no conflicts with other ranks in the same cloud. Enhanced Read Clouds reduce the risk of conflicting ranks and make it more likely that reads can be promoted.

We used Minerva to improve the specificity of short read taxonomic assignments obtained from Kraken, a popular pseudo-alignment based tool [18]. We selected Kraken because it was found to have good precision but relatively poor recall in a study by McIntyre et al. [13].

Using the technique described above we were able to rescue a large number of reads from unspecific taxonomic assignments. We rescued reads using both enhanced Read Clouds and standard Read Clouds. In every case rescue with enhanced Read Clouds matched or outperformed rescue with standard Read Clouds. All cases where rescue with enhanced Read Clouds outperformed standard Read Clouds for Dataset 1 are shown in table 3. All observed taxonomic assignments were correct after promotion. Without enhanced barcodes many annotations cannot be rescued or are incorrect.

**Table 3:** Taxonomic Promotion

| Original Rank | Promoted Rank | Enhanced | Standard | Difference | Ratio |
| --- | --- | --- | --- | --- | --- |
| Bacteria | *Enterobacter cloacae* | 3 | 2 | 1 | 1.5 |
| Proteobacteria | *Enterobacter cloacae* | 24 | 17 | 7 | 1.41 |
| Gammaproteobacteria | *Enterobacter cloacae* | 21 | 13 | 8 | 1.62 |
| Enterobacterales | *Enterobacter cloacae* | 87 | 72 | 15 | 1.21 |
| Enterobacteriaceae | *Enterobacter cloacae* | 765 | 642 | 123 | 1.19 |
| Bacteria | *Escherichia coli* | 9 | 6 | 3 | 1.5 |
| Proteobacteria | *Escherichia coli* | 8 | 7 | 1 | 1.14 |
| Enterobacterales | *Escherichia coli* | 17 | 13 | 4 | 1.31 |
| Enterobacteriaceae | *Escherichia coli* | 9221 | 7846 | 1375 | 1.18 |
| Escherichia | *Escherichia coli* | 201 | 198 | 3 | 1.02 |
| Gammaproteobacteria | *Pseudomonas antarctica* | 3 | 2 | 1 | 1.5 |
| Pseudomonas | *Pseudomonas antarctica* | 256 | 200 | 56 | 1.28 |

**Table 4:** The number of reads which could be promoted using standard or enhanced read clouds in a deconvolution of dataset 1. This figure uses the same deconvolution as figure 2. Cases where enhanced read clouds did not outperform standard read clouds are omitted, there are no cases where standard outperformed enhanced.

## 3 Discussion

We have introduced **Minerva** a graph based algorithm to provide a solution to the barcode deconvolution problem. By design Minerva provides conservative solutions to barcode deconvolution for metagenomics and uses essentially no information (except kmer overlaps) about the sequences being clustered. As such Minerva is a relatively pure demonstration of how information can be extracted from Linked-Reads. With some modification the algorithms underlying Minerva may even be useful for detecting structural variations and other genetic structures in the human genome.

However, the current version of Minerva could be enhanced by leveraging a number of practical sequence features: such as known taxonomic assignment, GC content, tetramer frequency, or motifs. These have been shown to be good indicators of lineage in metagenomics and could be easily incorporated to improve Minerva’s clusterings. In particular, taxonomic assignments could be incorporated into Minerva to evaluate barcode deconvolution, since there is no a-priori reason to think reads with a known taxonomic classification would be deconvolved more effectively than reads that could not be classified.

The current version of Minerva provides reasonable performance but still represents a potential bottleneck for workflows using Linked Reads. A large performance issue is Minerva’s routine to calculate the size of an intersection between two sets which is naive and exact. Jain *et al.* [9] has shown that bloom filters can be effectively used to speed up the calculation of set intersection in biology with acceptable errors. Future versions of Minerva could employ similar techniques to improve performance. Minerva uses the same parameters to process every barcode, however the nature of Linked-Read sequencing provides a rich source of information that could be used to optimize model parameters for deconvolving individual barcodes. This would require a more thorough mathematical model of linked reads which we leave to a future work. Similarly, external sequence annotation could be incorporated as a practical approach to setting parameters for individual barcodes though it is unlikely that such a technique would generalize to non-microbial applications.

Of particular interest to us is the possibility of using Minerva to directly improve downstream applications. For simple applications Minerva may be used with a single set of parameters to produce a deconvolution that meets certain requirements. For applications built to take advantage of barcode deconvolution Minerva could be run with multiple parameters to produce increasingly strict tiers of enhancement. This may be particularly important for de Bruijn graph assembly. DBG assembly typically relies on effectively trimming and finding paths through a de Bruijn graph. Multiple tiers of linkage between reads could be used to inform trimming or path finding programs about likely paths and spurious connections. This could likely be modeled either as an information theory or probabilistic approach depending on the situation and assembler.

Overall we believe that Minerva is an important step towards building techniques designed to take advantage of Linked-Reads. Linked-Reads have the potential to dramatically improve detection of large genetic structures without dramatically increasing sequencing costs and while taking advantage of existing techniques to process short reads.

## 4 Methods

We have developed a graph based algorithm to subdivide reads from the same Read Cloud into groups that, ideally, solve the barcode deconvolution problem.

The core intuition behind our approach is that reads from the same Read Cloud from the same fragment will tend to overlap with similar sets of reads from other Read Clouds. Critically, if the total genome length in a sample is large enough, a pair of Read Clouds is unlikely to contain reads from more one overlapping genomic region. We justify this statement in section 4.1.

### 4.1 Mathematical Justification of the Model

We have developed a simple model to justify our statement that reads from the same fragment will overlap with reads from similar sets of Read Clouds. This model is similar to empirical results and can be used to inform the parameters used for deconvolution.

First, we develop a model for drawing fragments of DNA from genomes in a metagenomic sample. For simplicity, We model each microbial genome *G_i_* in a metagenome *G* as a discrete collection of exactly *N_g_* fragments *F_i_. N_g_* is the same for all genomes. The probability of selecting a given fragment *F_i,j_* from a given microbial genome *G_i_* is given by a uniform distribution. We model the probability of selecting a given genome as a Geometric distribution; this choice is motivated by observations of real microbial communities that tend to be dominated by 1-2 species with a long tail of lower abundance species.

The probability of selecting a particular fragment *F_i,j_* given that we are drawing fragments from genome *G_i_* is

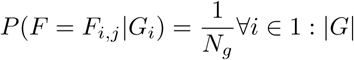

For simplicity we assume the abundance of genomes *G*_1_*,G*_2_, ⋯ is sorted in descending order by their index. The probability of selecting a given genome *G_i_* is

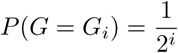

This gives us the probability of drawing a single given fragment *F_i,j_* without a given genome.

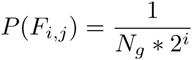

The probability that two fragments *F_w,x_, F_y,z_* are the same given that their genomes *G_w_, G_y_* are the same is

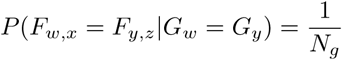

The probability that two genomes *G_w_*, *G_y_* are the same is

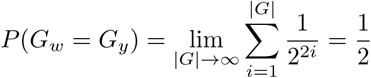

Let *p_f_* be the probability that two fragments *F_w,x_*, *F_y,z_* are the same without given genomes. We have

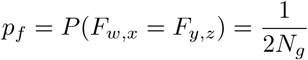

Second, we develop a generative model for assembling a Read Cloud from a set of fragments. We model each Read Cloud as a selection of *N_f_* fragments drawn from the set of all possible fragments. We refer to the set of fragments in a given Read Cloud as *R_i_* For simplicity we do handle the case where two Read Clouds both contain multiple fragments from the same class, this case is very unlikely with parameters relevant to our scenario (1 in 25,000 with the parameters given below).

Let *X*(*k*) be the probability that two Read Clouds *R_i_* and *R_j_*, both with *N_f_* fragments, share exactly *k* fragments. In other words any fragment in *R_i_* overlaps with at least one fragment in *R_j_* and vice versa.

We have

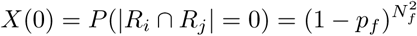

This is simply because none of the 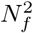 possible pairs of fragments (i.e. one in *R_i_* and one in *R_j_*) overlap.

We also have

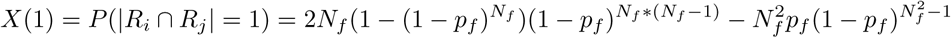

Here exactly one fragment in *R_i_* overlaps with one or more fragments in *R_j_* or vice versa. While it is extremely unlikely that we observe overlap of a fragment in *R_i_* with more than one fragment in *R_j_*, we handle this case in our equations because this is allowed in our approximate generative model. We have 2*N_f_* possibilities to select a fragment in either *R_i_* or *R_j_*. This fragment must overlap with at least one fragment in the other Read Cloud (i.e. the term 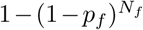). No other pair of fragments must overlap (i.e. the term 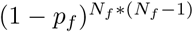) and because we double-counted cases where exactly one fragment in *R_i_* overlaps with exactly one in *R_j_* we subtracted the term 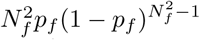.

The probability that two Read Clouds share more than one fragment is

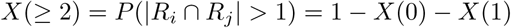

We choose reasonable, conservative (compared to our observed data), values for all parameters *N_f_* = 5, *N_g_* = 100, |*G*| = 10 and obtain the following estimates.

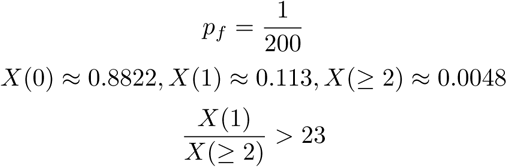

We find that it is about 23 times more likely to have exactly one overlapping fragment between two Read Clouds than multiple overlapping fragments in our mathematical model. We verified this through simulation and obtained a similar ratio of 1 to 40 (the discrepancy occurs because of how our simulation samples the geometric distribution). This is true even with conservative parameters chosen to minimize the ratio. This is important because it means we are likely to avoid a large number of spurious connections between genomic regions that could lead to poor deconvolution. However, this model does not account for the fact that individual fragments may have similar sequences, which is a major source of noise for **Minerva**. To reduce this noise, we use the parameters of this model to justify removing any overlaps that occur far more often than expected.

On average, each fragment in a dataset is only fractionally covered at a rate of *C_r_* (with a read depth of 1). While the precise coverage might vary between fragments this parameter can be used to estimate the size of overlaps between fragments and their expected sparsity. Two long fragments would be expected to overlap at 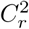 points in their overlap. In cases where fragments overlap much more frequently than 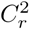 over their lengths it can be inferred that the sequence present is too repetitive or common to be useful for deconvolution.

These facts are used in **Minerva** to filter connections between repetitive regions, restrict overlaps to regions of a certain length, and to heuristically filter comparisons between Read Clouds unlikely to have significant overlap. This carries practical performance benefits and reduces errors.

### 4.2 A Graph Based Algorithm for Barcode Deconvolution

#### 4.2.1 Summary

We have developed a graph based algorithm that effectively deconvolves the reads within a given Read Clouds. The model constructs a bipartite graph between all reads with a given Read Cloud and all other Read Clouds. Reads have an edge to a Read Cloud if they are found to contain a kmer that is specific to exactly one read in the foreign Read Cloud. Once the bipartite graph is constructed Read Clouds and reads with too many or too few edges (by user supplied parameters) are removed. The filtered bipartite graph is used to construct an adjacency matrix between reads and the matrix is clustered into groups of reads. This algorithm is *O(n*^2^) over the number of Read Clouds, though we note that the number of Read Clouds is a constant for each specific technology that could be used.

The specific steps in our algorithm are as follows:

1. Read Clouds are parsed, Read Clouds below a certain size (dropout) are dropped
2. Each read in each Read Cloud is parsed into a set of minimum sparse kmers
3. Each Read Cloud above a certain size (anchor dropout) is compared to all other read clouds. The read cloud being compared is called the ‘anchor’
4. A bipartite graph is constructed between the reads in the anchor and all other read clouds based on kmer overlap
5. The bipartite graph is reduced to a graph between reads in the anchor
6. The read graph is broken into discrete clusters which are output as solutions to the barcode deconvolution problem

#### 4.2.2 The Model

Initially each Read Cloud in a given dataset is parsed into a set of minimizing kmers (see section 4.4, figure 4 part 2). Global counts for kmers are retained. Once parsing is complete, kmers that occur exactly once or many times more than the average (10 times more, by default) are discarded. Singleton kmers cannot occur in more than one barcode and kmers that are too common tend to create false positives (these kmers appear to originate from low complexity or conserved regions). This process is analogous to removing stop words in Natural Language Processing applications. A map of kmers to reads is retained for each Read Cloud.

**Figure 4:**
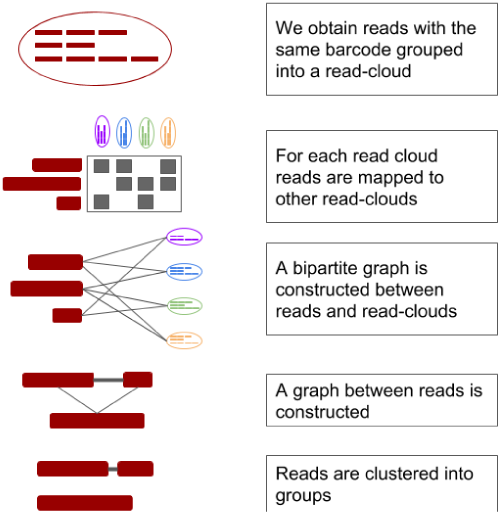
From top 1) Fragments are generated 2) Fragments are sequenced and tagged 3) Reads in a given barcode are aligned to other barcodes 4) A bipartite graph between reads and barcodes is constructed 5) A graph between reads that co-occur with barcode is constructed 6) Reads are clustered into groups

After parsing, the set of reads in a given Read Cloud is compared to every other Read Cloud (figure 4 parts 3 and 4). Comparisons between Read Clouds that share too many kmers are discarded as these likely represent low complexity or evolutionarily conserved regions as opposed to real overlaps. Comparisons between Read Clouds that share too few kmers are also rejected to improve performance. The intersection of the kmer sets between the given Read Clouds and all Read Clouds that passed filtering is calculated.

A bipartite graph is constructed by creating nodes for every read in the Read Cloud being processed and every Read Cloud that was not filtered out. Edges are only added between read-nodes (left nodes) and Read Cloud-nodes (right nodes). An edge is drawn between a read-node and a Read Cloud-node if, and only if, the read shares a kmer with the given Read Cloud. This is a fast proxy measure for read overlap. Finally, any Read Cloud-node with degree above a given threshold is discarded.

Each bipartite graph representing the reads from a given Read Cloud is given a final round of filtering where reads that matched too many foreign Read Clouds are removed based on a user supplied threshold. The filtered bipartite graph is converted to an adjacency matrix of reads where the similarity between reads is equivalent to the number of Read Clouds that both reads overlapped with (figure 4 part 5). This adjacency matrix is converted to a binary matrix by setting all values below a user supplied threshold to zero and all remaining values to one (figure 4 part 6).

All connected components in the binary matrix are found. Connected components consisting of single reads are discarded, the remaining components define clusters. This process is analogous to DBSCAN [4] for graphs.

### 4.3 Information Theory Bounds on Barcode Deconvolution

We note that the barcode deconvolution problem on the graph based model we describe in section 4.2 is analogous to the community recovery problem [6] in Information Theory. In particular, 3’ barcodes provide linkage information between pairs of reads. We use this linkage information to construct a graph between the reads being deconvolved with the expectation that reads from the same fragment will have a better chance of being linked than reads from different fragments. Formally we say that two reads are connected with probability *p* if they are from the same fragment and probability *q* if they are from different fragments. Termed differently *p* is the true positive rate while *q* is the false positive rate.

For clarity we note that this model is distinct from the model we developed previously to justify why overlaps between read clouds were likely to be useful for deconvolution.

If we make a simplifying assumption that all fragments in our Read Cloud produce equal numbers of reads we can use the formula determined by [8] to determine the minimum connectivity of linking 3’ barcodes necessary to deconvolve our reads. We define the number of reads per fragment as 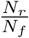, where *N_r_* is the total number of reads in a Read Cloud and *N_f_* is the number of fragments in the given Read Cloud.

For the community recovery problem [8] have provided a lower bound on the size of graph that can be accurately clustered given values of *p* and *q*, regardless of the algorithm used. If a graph is smaller than this threshold it is unlikely that it will be possible to distinguish clusters from spurious edges. This boundary requires us to assume that all fragments with the same 3’ barcode produced equal numbers of reads. Using the definitions above this definition we can apply the following inequality to Read Cloud deconvolution:

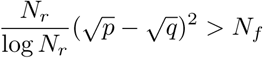

Using the model developed in 4.1 and a simulation we estimate the maximum true positive rate *p* to be 0.998 and we estimate the minimum false positive rate *q* to be 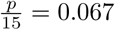. We note that these values do not account for multiple sources of error, notably sequence homology, and should be interpreted as a best case scenario. Using these values we can reduce the previous equation:

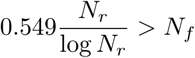

If a barcode deconvolution graph does not meet this inequality it is unlikely that it will be possible to accurately reconstruct all clusters. More generally this formula can be used to estimate the minimum number of reads and maximum error rates which can lead to effective barcode deconvolutions. In principle, this inequality should apply to all barocode deconvolution algorithms that can be formulated as a graph. However different algorithms may have different values of *p* and *q*. We also note that the above formula is based on asymptotic behavior for graphs with thousands of nodes. We observed that typical deconvolution graphs in our model have fewer than 50 nodes.

### 4.4 Minimum Sparse Hashing

Minerva frequently tests whether pairs of reads overlap. Many solutions to finding overlaps between reads exist, such as: sequence clustering algorithms, sequence aligners, and kmer matching. These techniques typically make trade offs between overall performance and error rates. Since Minerva is meant to be relatively fast and can tolerate some errors we elected to use a minimal sparse hash of kmers to match read pairs. This technique reduces the number of unique kmers Minerva uses to find overlaps which reduces runtime and RAM usage.

Minimum sparse hashing was originally developed independently for biological sequence search and natural language document search [10, 16] (in natural language search the technique is referred to as winnowing). While the original application of this technique in biology defined minimization as the lexicographic minimum of a set of sequences we use a uniform random hash function to determine the minimal sequence in a set. This is a common practical enhancement recently detailed by [15].

Minimum sparse hashing for sequences takes three parameters, a length *k,* a window size *w*, and a hash function *h*. Given a set *k* of *n*, *n* ≥ *w* kmers, the min-sparse hash computes the hash *h* of each kmer then selects the kmer with the smallest numerical hash from each consecutive set of *w* kmers in *K*. The final set of minimizers is the unique set of kmers generated, *W*. Each consecutive window shares *w* − 1 kmers so there is a good chance that each window shares the same minimum with its predecessor. Formally 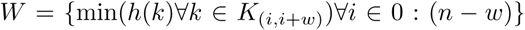. This algorithm guarantees that any pair of reads with an exact overlap of at least *w* + *k* − 1 bases will share at least one minimum sparse kmer while drastically reducing the number of kmers which must be stored in memory (figure 5). In certain implementations minimum sparse kmers may also improve performance by allowing a kmer that can be stored in a single 64 bit cell (*k* ≤ 32) of memory to represent a longer sequence.

**Figure 5:**
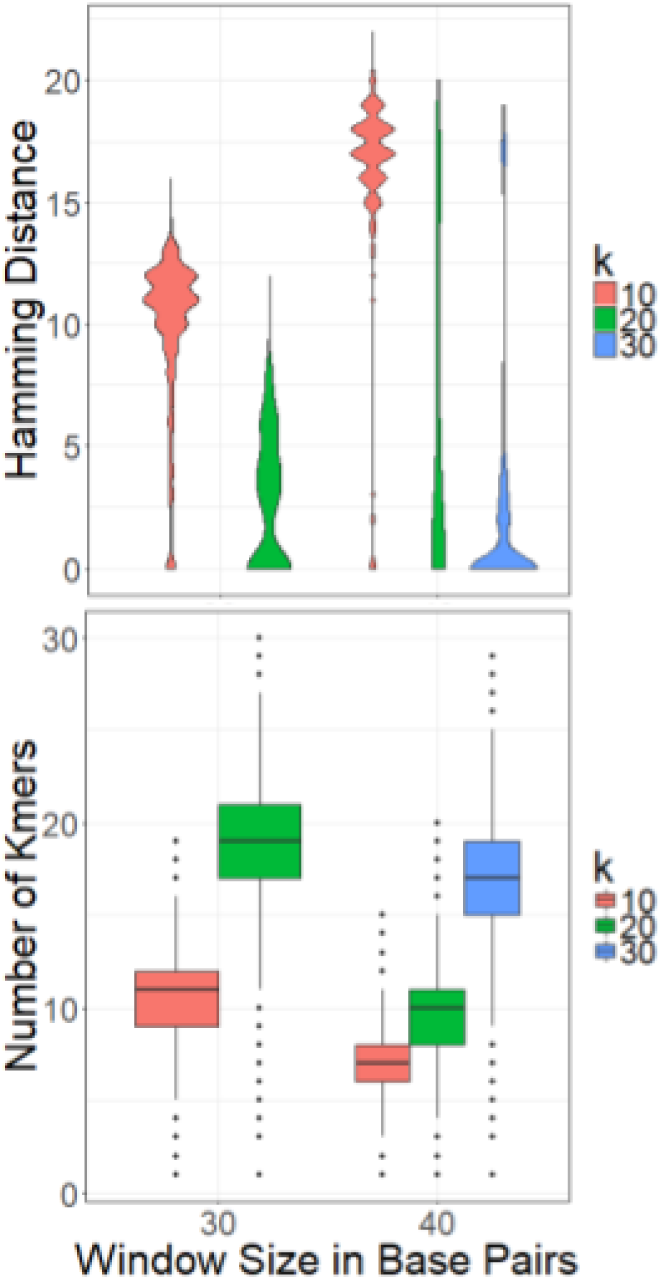
Top, Hamming distance between windows that share minimizing kmers, using various parameters. Bottom, number of representative minimizing kmers per read

Minimum sparse kmers are prone to false positives when presented with similar, but not identical, runs of *w* bases in read pairs. We measured this phenomenon by comparing all kmers of length *w* from pairs of reads that share a minimum sparse hash. Figure 5 shows the minimum hamming distance for windows of length *w* between reads that share a minsparse hash. When *k* is larger the average hamming distance is smaller though outliers persist. Small values of *k* produce many distant false positives. Raising *k* from 20 to 30 (*w* = 40) improved accuracy and precision to the point where false positives could be controlled using downstream techniques.

The mathematics that underly minimum sparse hashing may also be used to efficiently approximate the overlap between sets, another important operation for Minerva. We did not use this technique in our current implementation of Minerva but plan to explore this for later versions.

## 5 Data Access

The version of Minerva used to write this paper is freely available at https://github.com/dcdanko/minerva_barcode_deconvolution. The data sets used to evaluate Minerva are also available. The download links can be found on the same Github page.

## 6 Acknowledgement

We acknowledge Victoria Popic and her team at Illumina Inc. for fruitful discussions on the problem and her help in benchmarking. We sincerely thank 10X Genomics Inc. for providing us valuable real sequencing data sets used in this study for evaluations. We also thank Stephen Williams of 10X Genomics Inc. for coordinating the 10X Metagenomics consortium in which our team participates. DCD and DM were supported by the Tri-Institutional Training Program in Computational Biology and Medicine (CBM) funded by the NIH grant 1T32GM083937. This work was also supported by start-up funds (Weill Cornell Medicine) to Iman Hajirasouliha.

## 7 Supplementary Materials

### 7.1 Read Clouds

We show a histogram of observed Read Cloud sizes in figure 6. This histogram is truncated and does not show a long tail of large Read Clouds. This figure shows that the majority of Read Clouds are small.

**Figure 6:**
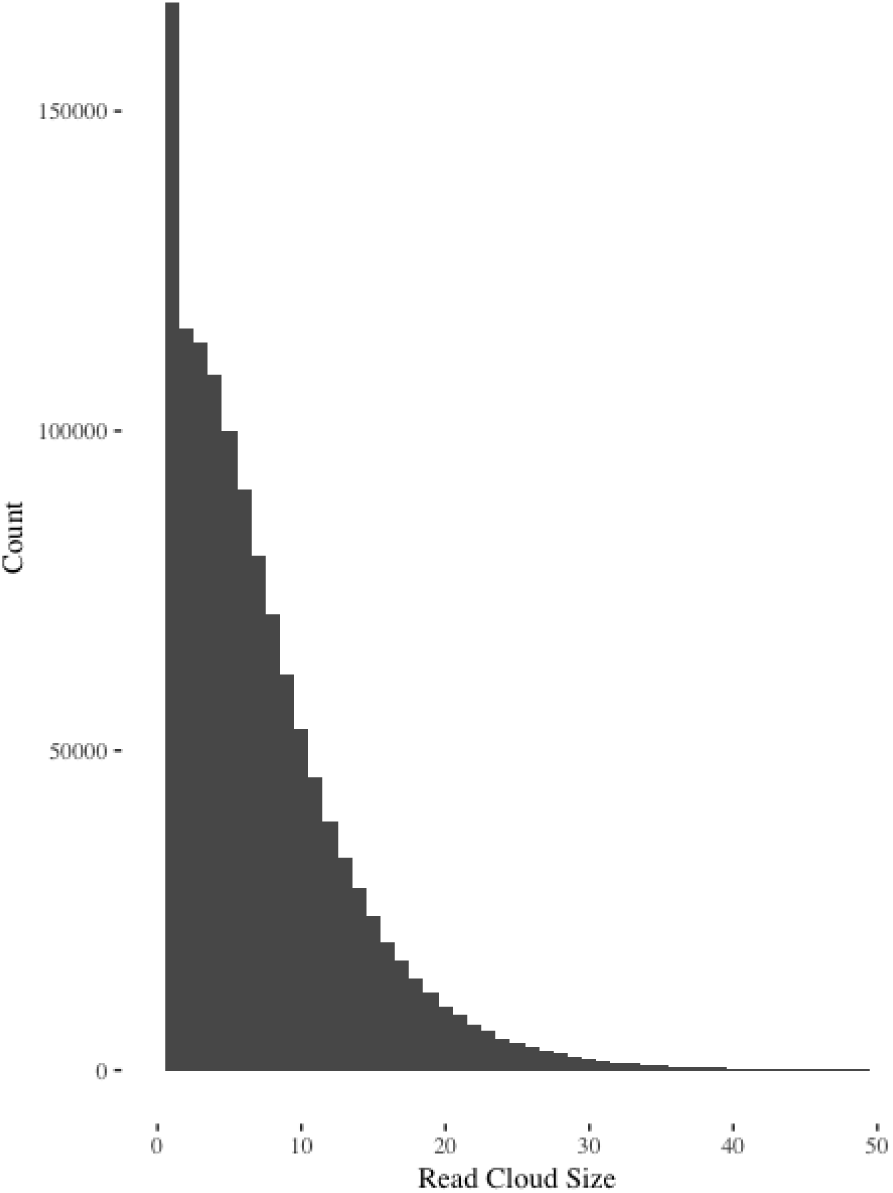
Histogram of Read Cloud Sizes in Dataset 1

**Figure 7:**
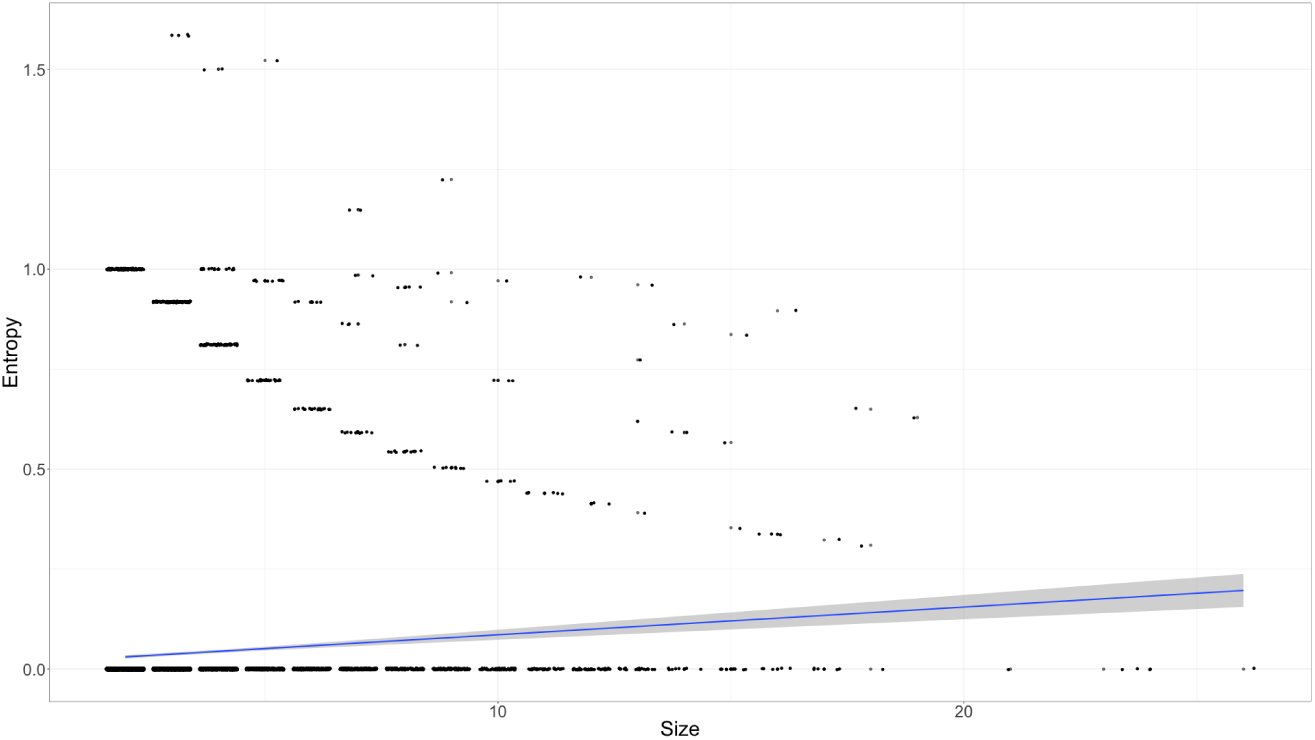
Entropy of enhanced Read Clouds only increases slightly with size. Jitter added to points for clarity.

We show a scatter plot of enhanced Read Cloud size versus entropy. Entropy only increases slightly with enhanced Read Cloud size.

### 7.2 Effect of Parameters

We summarize the impact of various parameters on model performance in figure 8.

**Figure 8:**
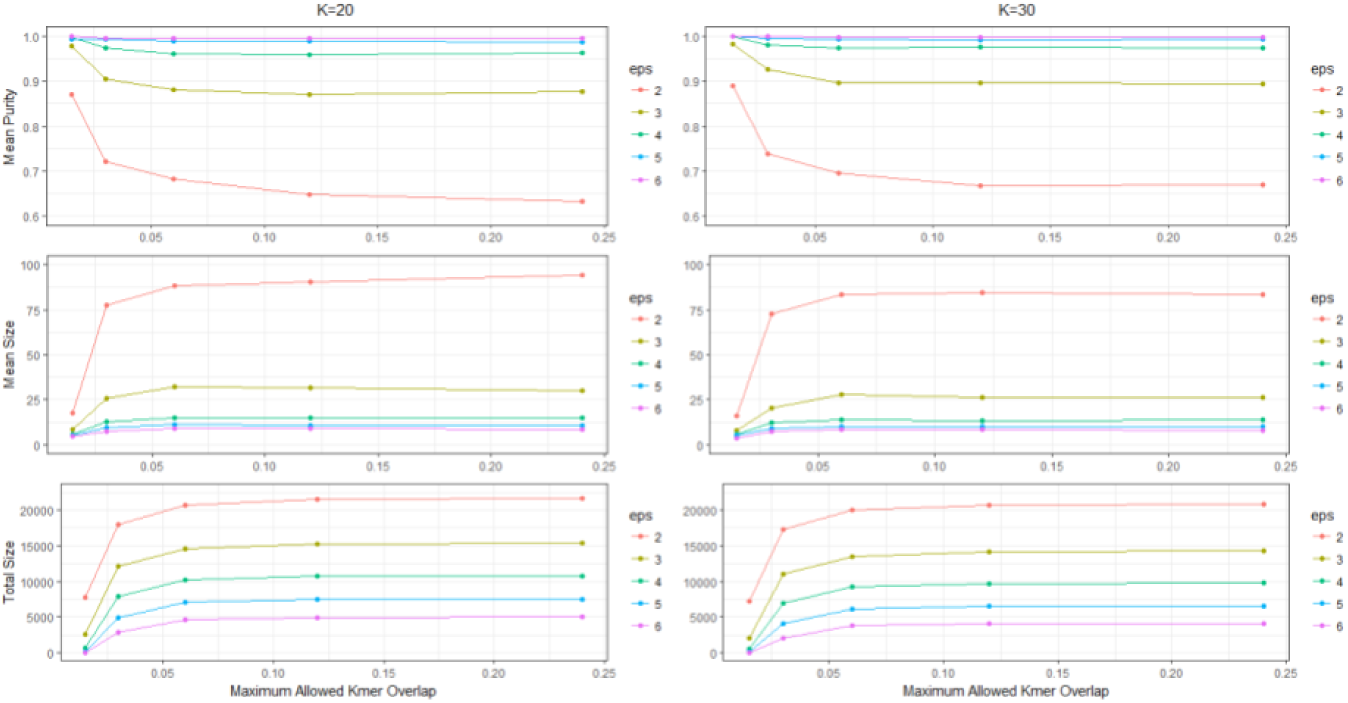
Variation of parameters

### 7.3 Enhancements to our Algorithm

Depending on the parameters and dataset used the clustering step may produce clusters that are large and imprecise or small and incomplete; this is partly a consequence of using a small number of discrete components to define connections. To mitigate this issue we have introduced two enhancements to the clustering step that may optionally be used. We have based these enhancements on the two step procedure used for community recovery by [2] and [14].

We term the first enhancement read-rescue; in some instances it is useful to rescue components that consist of single reads by connecting them to clusters that they unambiguously share connections with. To do this all single reads are checked for connections to any read in a cluster. If the single read is connected to enough reads (by a user supplied parameter) in no more than one cluster the read is added to that cluster. We term the second enhancement edge-cracking; when clusters are too large it is useful to break certain edges that bridge two individually well-connected components. To do this all edges in the adjacency matrix are considered. If the two reads attached to an edge do not share enough neighboring reads (by a user supplied parameter) the edge is removed from the adjacency graph.

### 7.4 Failed Models

While developing this model we tested several variants that performed poorly. In this section we report these models as negative results, parameters and outcomes may be found in the supplement. We define three distinct components of our model:

1. Barcode reduction, a technique to generate a matrix with rows representing reads and columns related to barcodes
2. Cell weighting, a technique to reweight the cells of our reduced matrix
3. Clustering, alternate techniques to cluster our reduced, reweighted matrix.

We tried several techniques to reduce our matrix (our final technique was to build a bipartite graph that was converted into an adjacency matrix) including: Principal Component Analysis, Latent Dirichlet Allocation (treating each read as a document of barcodes), the jaccard, dice and kulsinski distance matrices between reads.

For most reductions we tried to reweight our matrices by normalizing row and column sums. We also tried term frequency - inverse document frequency and just inverse document frequency. For our final model we did not use any reweighting scheme.

To cluster our reduced and reweighted matrix we tried xmeans, kmedoids, hierarchical clustering with various linkages and Markov clustering. Hierarchical clustering yielded reasonable performance but dbscan was faster, had more intuitive parameters, and had slightly better performance.

Additionally, we tried to use LDA globally (treating each barcode as a document of kmers) to discover sets of kmers that uniquely identified fragments. This technique provided surprisingly good barcode deconvolution (and automatically defined globally clusterings of barcodes) but was extremely computationally costly and did not yield results as good as our final model.

